# Hardy Weinberg Exact Test In Large Scale Variant Calling Quality Control

**DOI:** 10.1101/095521

**Authors:** Zhuoyi Huang, Navin Rustagi, Degui Zhi, L. Adrienne Cupples, Richard Gibbs, Eric Boerwinkle, Fuli Yu

## Abstract

Hardy Weinberg Equilibrium (HWE) test is widely used as a quality control measure to detect sequencing artifacts like mismapping, allelic dropout and biases. However, in the high throughput sequencing era, where the sample size is beyond a thousand scale, the utility of HWE test in reducing the false positive rate remains unclear. In this paper, we demonstrate that HWE test has limited power in identifying sequencing artifacts when the variant allele frequency is lower than 1% in a variant call set produced from more than five thousand whole genome sequenced samples from two homogeneous populations. We develop a novel strategy of implementing HWE filtering in which we incorporate site frequency spectrum information and determine the p-value cutoff which optimizes the tradeoff between sensitivity and specificity. The novel strategy is shown to outperform the exact test of HWE with an empirical constant p-value cutoff regardless of the sequencing sample size. We also present best practice recommendations for identifying possible sources of false positives from large sequencing datasets based on an analysis of intrinsic biases in the variant calling process. Our novel strategy of determining the HWE test p-value cutoff and applying the test to the common variants provides a practical approach for the variant level quality controls in the upcoming sequencing projects with tens to hundreds of thousand of samples.

## Introduction

The decreasing cost of sequencing makes it possible to sequence large cohorts with moderate to low coverage at million sample size scale in the coming years. An analysis of variant callsets produced from large sequencing datasets are also presenting an unprecedented resolution at low frequency variants [1]. The increased role of rare variants in common diseases has also been documented [2], and improving the yield of rare variants in low coverage whole genome datasets has become extremely important. To increase the yield of high fidelity variants, quality control of variant call sets has been a crucial step in the past large scale projects [3, 4]. More specifically, low sensitivity in variant site discovery and low specificity due to false calling can reduce the discovery power of novel and rare variants in low coverage sequencing projects with a large sample size and median coverage less than 10-fold. However quality assessment and quality control (QA/QC) steps often rely on the estimation of the site frequency spectrum (e.g. allele frequency and genotype frequency) which in themselves are challenging in the low frequency range due to low sequencing coverage [5, 6].

As a variant site level quality control method, the exact test of Hardy Weinberg Equilibrium (HWE) has been widely used [7] to filter false positive sites which significantly deviate from the random mating hypothesis [4, 8]. Statistical approaches based on deviations from Hardy Weinberg Equilibrium have been frequently used to ascertain genotyping errors, and population stratification in case control studies. The Hardy Weinberg test has a long history in population genetics [9, 10] but the exact test for Hardy-Weinberg has recently gained prominence as a preferred QA/QC method for Genome Wide Association Studies(GWAS) [8, 11, 12]. The test has been particularly useful in scenarios when highly polymorphic sites are targeted, as in SNP array, across several hundred of samples in the same population. Even though a naïve application of HWE filter can result in incorrect inferences for SNP array data [13], HWE test based filters are extensively used in the past, including on datasets consisting of high coverage (>50x) Exome data [14]. While inbreeding events within a population can result in lower prevalence of heterozygotes, significant excess of heterozygotes in a homogeneous population is usually an indicator of sequencing and calling artifacts, such as sequencing errors, misalignment, low coverage allelic dropout, false calling or false imputation [7, 8].. Empirically choosing a constant HWE test p-value between 10^−2^ to 10^−6^ based on past large scale projects [4, 14], regardless of the sample size, population ethnicity and the variant allele frequency, makes it hard to compare the quality and integrate the variant call sets from different sequencing projects. Many standard population genetics tests for estimating important information like diversity, selection and demographic history are direct applications of site frequency spectrum information. The resolution of discovering low frequency variation in populations is unprecedented due to the increasing capacity of sequencing extremely large cohorts, thereby making it intuitive to incorporate allele frequency information into QA/QC procedure.

In this paper we present a unified picture of QA/QC procedure in which we systematically determine a HWE p-value cutoff optimizing the power of HWE test in different ranges of allele frequency spectrum for datasets with large sequencing sample size. Firstly, we analyze the performance of HWE test in identifying sequencing artifacts at different allele frequency ranges, in a low coverage sequencing project consisting of several thousands of samples from two homogeneous populations. We specially emphasize on the challenges in identifying sources of error using the HWE test at the rare end of the site frequency spectrum. Secondly, we propose a novel strategy of applying HWE test in an effective allele frequency range, with a p-value cutoff which optimizes the overall sensitivity and specificity. The novel strategy is shown to outperform the HWE test with an empirical constant p-value cutoff independent of the sample size. Thirdly, based on our experiences with an extremely large sequencing cohort, we present guidelines for designing a QA/QC procedure which combines the HWE exact test with site frequency information for increased sensitivity and specificity in variant callsets from future large scale studies.

## Results

The QA/QC analysis presented in this paper has been carried out on the SNV callset of the CHARGES Freeze 3 whole genome dataset (see Methods for more details). This dataset has 3396 European Americans (EuAm) samples and 1901 African Americans (AfAm) samples. The gold standard data consists of SNP array genotype information of a subset of 1850 European American samples within the Charge Freeze 3 dataset. Unless otherwise mentioned explicitly, all the results presented in this paper is for the 1850 EuAm samples in the CHARGE dataset for which there is corresponding SNP Array data.

### Hardy Weinberg Equilibrium test has limited power for low allele frequency range (*f*<1%)

For large cohort sizes (>1000), Hardy Weinberg Equilibrium (HWE) test has very limited power in filtering out false positives in the rare end of the frequency spectrum ( allele frequency *f*<1%). Comparing the variants called in the low coverage whole genome sequencing (WGS) in 1850 EuAm samples to the SNP array data, the alternate allele frequency of identified false positive sites in the WGS call set follows a bimodal distribution, aggregating at both the common end (allele frequency (*f*>10%) and the rare end (*f*<1%) of the allele frequency spectrum (red, yellow and grey bins in Figure 1). By applying the exact test of HWE to both the true positive and false positive sites with the p-value cutoff(*p*)>10^−4^. most common false positives are filtered (grey bins in Figure 1) out. Increasing the p-value cutoff does not result in significant reduction of rare false positives any further. Even for p-value cutoff *p*>0.1. a reduction in false positive sites with *f*<1% is not observed (red bins in Figure 1). while more common true positive sites are filtered (the difference between green and blue curves in Figure 1) out. This suggests that the exact test of HWE should be applied to the common variant sites with allele frequency above a critical limit, denoted as *f*_c_. However, filtering mechanisms need to model sequencing artifacts to get higher quality variants at the rare end of the frequency spectrum, as is discussed in more detail in the subsequent paragraphs.

**Figure 1:**
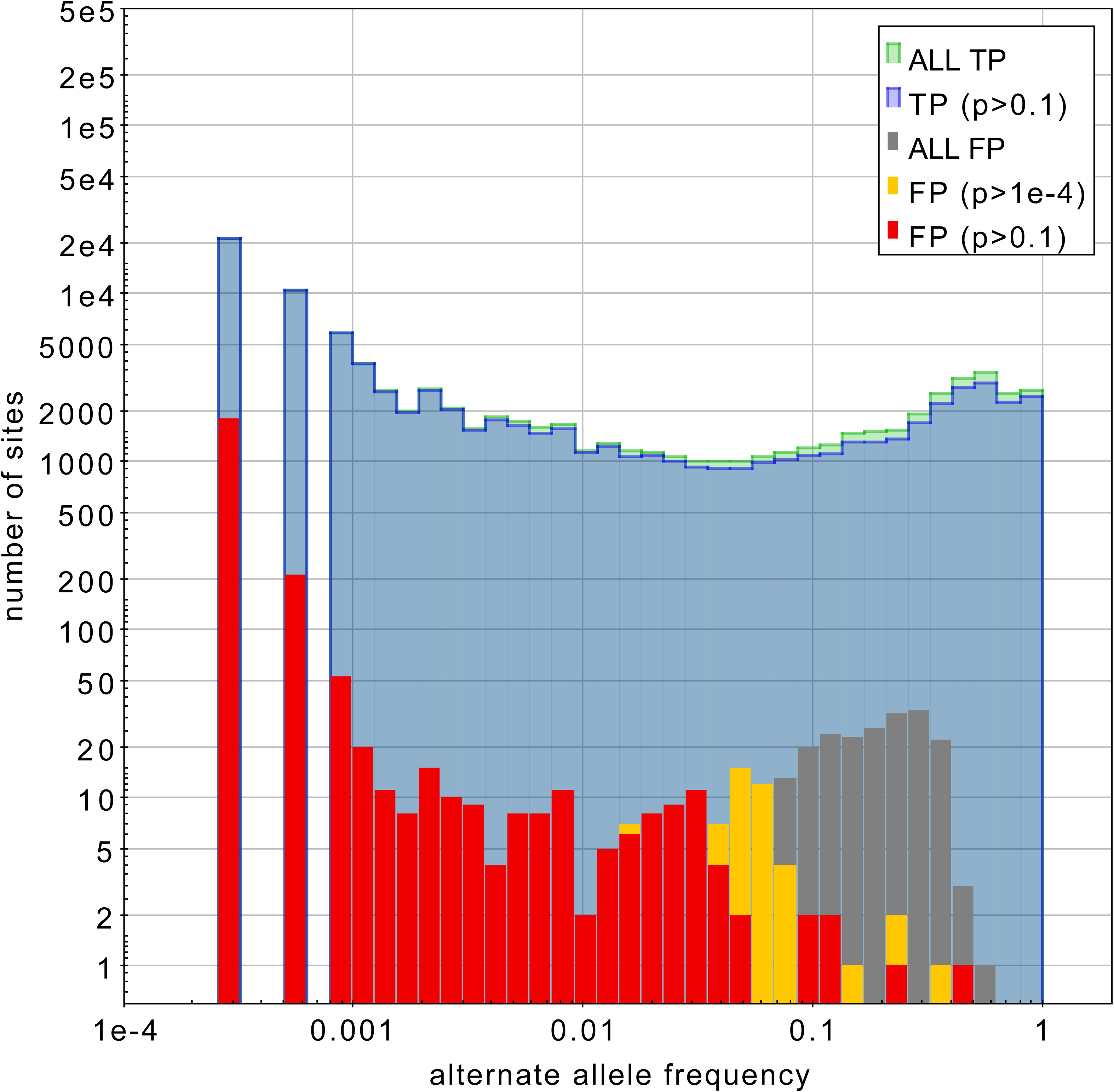
Alternate allele frequency distribution of true positive and false positive variant sites in the CHARGE European American samples taking SNP array data as gold standard. A subset of true positive sites with HWE test p-value cutoff *p*>0.1 is also shown (blue curve), while for false positive sites, subsets with *p*>0.1 (red bins) and *p*>10^−4^ (yellow bins) are overlaid on top of all false positive distributions (grey bins).

### Distribution of HWE p-values for false positive sites in the allele frequency

The true positive and false positive sites follow different distributions in the alternate allele frequency and HWE p-value (*f*-*p*) diagram (Figure 2). In the common allele frequency range (*f*>5%). where HWE test is effective, the p-value of most false positive sites (*p*<10^−4^) is significantly lower than the p-value of true positive sites (*p*>10^−3^). In the range *f*<1%, most false positive sites and true positive sites have indistinguishable p-values in the HWE test (green and red dots with *p*=1 in Figure 2).

**Figure 2:**
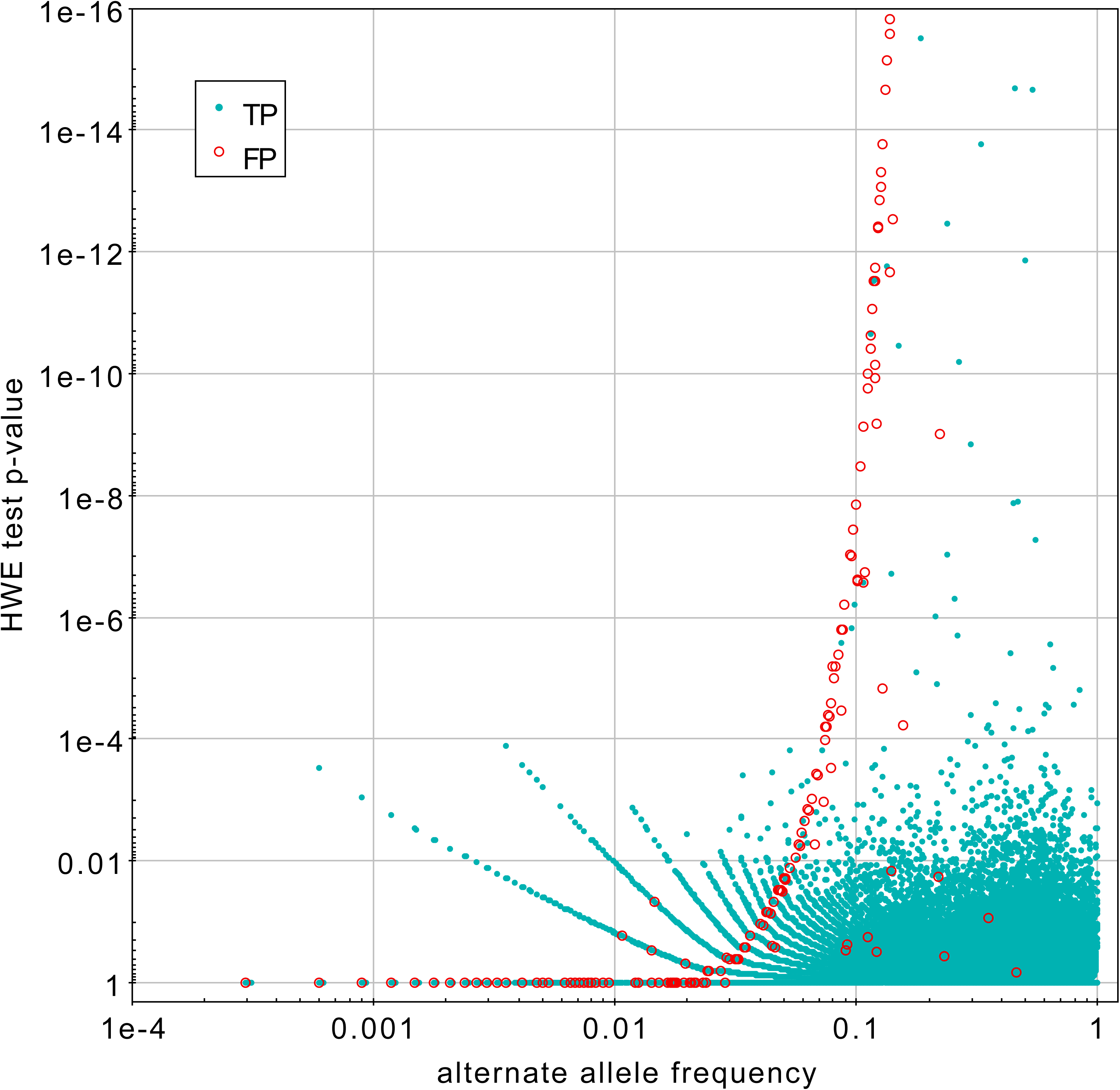
The distribution of true positive sites (green dots) and false positive sites (red circles) in the diagram of alternate allele frequency and HWE test p-value.

The variant sites on the false positive trajectory in the *f*-*p* diagram correspond to sites with lower number of homozygous Alt/Alt samples (*N_aa_*≤3 in Figure 2, see also Supplementary Figure 4). In common allele frequency range (alternate allele frequency *f*>5%), the extreme deficit of homozygous Alt/Alt samples indicates a significant excess of heterozygous Ref/Alt samples, which causes significant deviation from HWE. On the contrary, in the rare alternate low allele frequency range (*f*<1%), the deficit of homozygotes Alt/Alt is in agreement with HWE and the test is not informative to distinguish false positive from true positive sites, which may both have excess of heterozygotes. However the effect of sample sizes is reflected in the HWE p value plots as the false positive trajectory shifts towards the lower frequency end with increasing sizes (see Supplementary Figure 4). Other sequencing artifacts like mismapping and allelic dropout can also confound the HWE test and their role in developing effective QA/QC strategies is discussed below.

### The cause of false positive sites

To study the origin of false positive trajectory in the *f*-*p* diagram, we take the variant sites called in the CHARGE dataset with SNP array genotypes in a subset of shared samples, and compare the alternate allele frequency and HWE p-value (*f*,*p*) of WGS with that of genotypes in the SNP array (*f*_G_, *P*_G_). We investigated the sites with a significant (twice or greater) excess or deficit of alternate alleles in the *f*-*p* diagram (Figure 3). The sites with deficient alternate alleles *f*_G_>2*f* (Figure 3a) aggregate in the low allele frequency range (*f*<5%) with a generally high p-value *p*≥0.01 (see also histogram in Supplementary Figure 7). These are the sites with one or both of the alternate alleles dropped-out due to low sequencing coverage. On the other hand, sites with excess of alternate alleles, *f*>2*f*_G_ (Figure 3b), fall in two regions. In the common allele frequency range (*f*>5%), they are located at the false positive trajectory in *f*-*p* diagram, deviating clearly from the HWE. The large offset in the *f*-*p* diagram is contributed by possible misalignment and sequencing errors (see Supplementary Figure 9-10), which results in excess of heterozygotes, or contributed by false imputation, which introduces false alternate allele at sites with no coverage. The second region where sites with excess of alternate allele aggregate is the low allele frequency range, where the cause of offset is the same but high p-values in HWE test alone is not effective in separating them from the sites with deficit of alternate alleles (as in Figure 3a).

**Figure 3:**
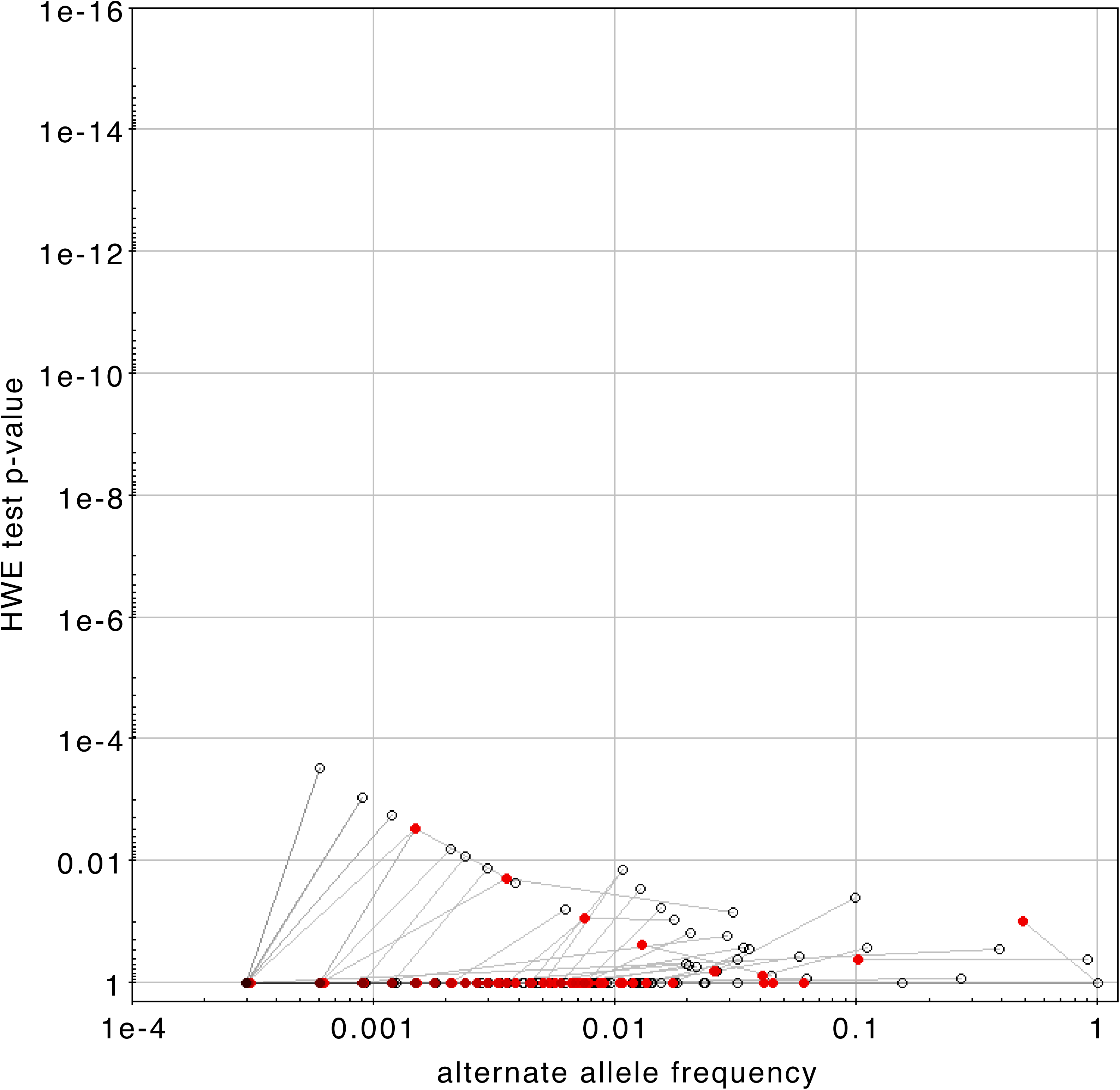
The difference of the allele frequency and HWE p-value between the WGS call set (filled points) and the SNP array data (open circles), at variant sites with deficit of alternate alleles (3a) and variants sites with excess of alternate alleles (3b).

We further investigate the genotyping error at sites with large offsets in *f*-*p* diagram. The overall genotype concordance at the true positive sites is high, on average (99.957%, 98.063%, 98.451%) for three genotypes (Ref/Ref, Ref/Alt, Alt/Alt) (Supplementary Table 1). At sites where variant allele are correctly called but per sample genotyping error causes excess of alternate alleles (0.92% of all true positive sites), the average proportion of genotypes (Ref/Ref, Ref/Alt, Alt/Alt) in SNP array data is (99.887%, 0.090%,0.023%), while in the WGS call set, it is (99.463%, 0.535%,0.002%) (Table 1). At these sites, 90% of the Alt/Alt samples and some Ref/Ref samples are incorrectly genotyped as Ref/Alt. On the other hand, for sites with deficit of alternate alleles (5.14% of all true positive sites), the average proportion of genotypes of both Ref/Alt and Alt/Alt drops from 0.255% and 0.262%, in the SNP array, to 0.110% and 0.006%, respectively, in the low coverage WGS call set (Table 1), as a result of low sequencing coverage.

**Table 1.** The difference in the average genotype proportion of the variant sites between SNP array and WGS call set.

|  | All TPs |  |  | TPs with excess of Alt |  |  | TPs with deficit of Alt |  |  |
| --- | --- | --- | --- | --- | --- | --- | --- | --- | --- |
|  | Ref/Ref | Ref/Alt | Alt/Alt | Ref/Ref | Ref/Alt | Alt/Alt | Ref/Ref | Ref/Alt | Alt/Alt |
| SNP array | 88.34% | 10.08% | 6.57% | 99.89% | 0.09% | 0.02% | 99.48% | 0.26% | 0.26% |
| Called | 83.51% | 9.97% | 6.52% | 99.46% | 0.54% | 0.00% | 99.89% | 0.11% | 0.01% |

At false positive sites, besides the genotyping error as mentioned above, all samples are genotyped against as alternate allele which is falsely called due to similar reasons. We find that most of the sites with these genotyping errors are rare with *f*<1% (Supplementary Figure 7), is consistent with the excess of false positive sites in the low frequency range where the HWE filtering power is limited.

### The strategy to determine optimal p-value cutoff

Within the allele frequency range where the HWE test is informative, we can determine the optimal p-value cutoff. We take the difference between the true positive rate (TPR) and false positive rate (FPR) as *δ*=TPR-FPR. It is an evaluator of the HWE filtering performance, similar to the receiver operating characteristic (ROC) analysis. Since the 1+*δ* is the sum of sensitivity and specificity, we define the cutoff which maximizes *δ* as the optimal tradeoff between the two. Applying HWE to the full allele frequency spectrum only gives maximum *δ*=0.08 at p-value cutoff *p*>10^−3^ (Figure 4 red dash line). Since the p-value of true positive and false positive sites is as high as 1 below certain allele frequency, we can choose it as a limiting allele frequency *f*_c_, and only apply HWE test to the common sites with alternate allele frequency *f*>*f*_c_ (see Methods). For instance, when the sample size is 1850, and the number of homozygotes *N*_aa_=0, the limiting allele frequency is *f*_c_=0.0239. Above this allele frequency, the optimal p-value cutoff gives *δ*=0.8 (Figure 4 blue dash line), which is increased by 10-fold compared to a constant p-value applied to all allele frequency spectrum. The filtering performance is significantly improved due to two factors. First, when *f* > *f*_c_, the true positives and false positives follow very different p-value distribution as is mentioned above. Second, towards the low allele frequency end, where the HWE test is non-informative, the number of variant sites is exponentially increasing, therefore excluding the low frequency range in the test significantly increases the performance. With the optimal p-value cutoff, the overall sensitivity, specificity and FDR of the WGS call set is 85.709%, 97.727% and 2.388%, respectively, taking SNP array data as the gold standard (Supplementary Table 1).

**Figure 4:**
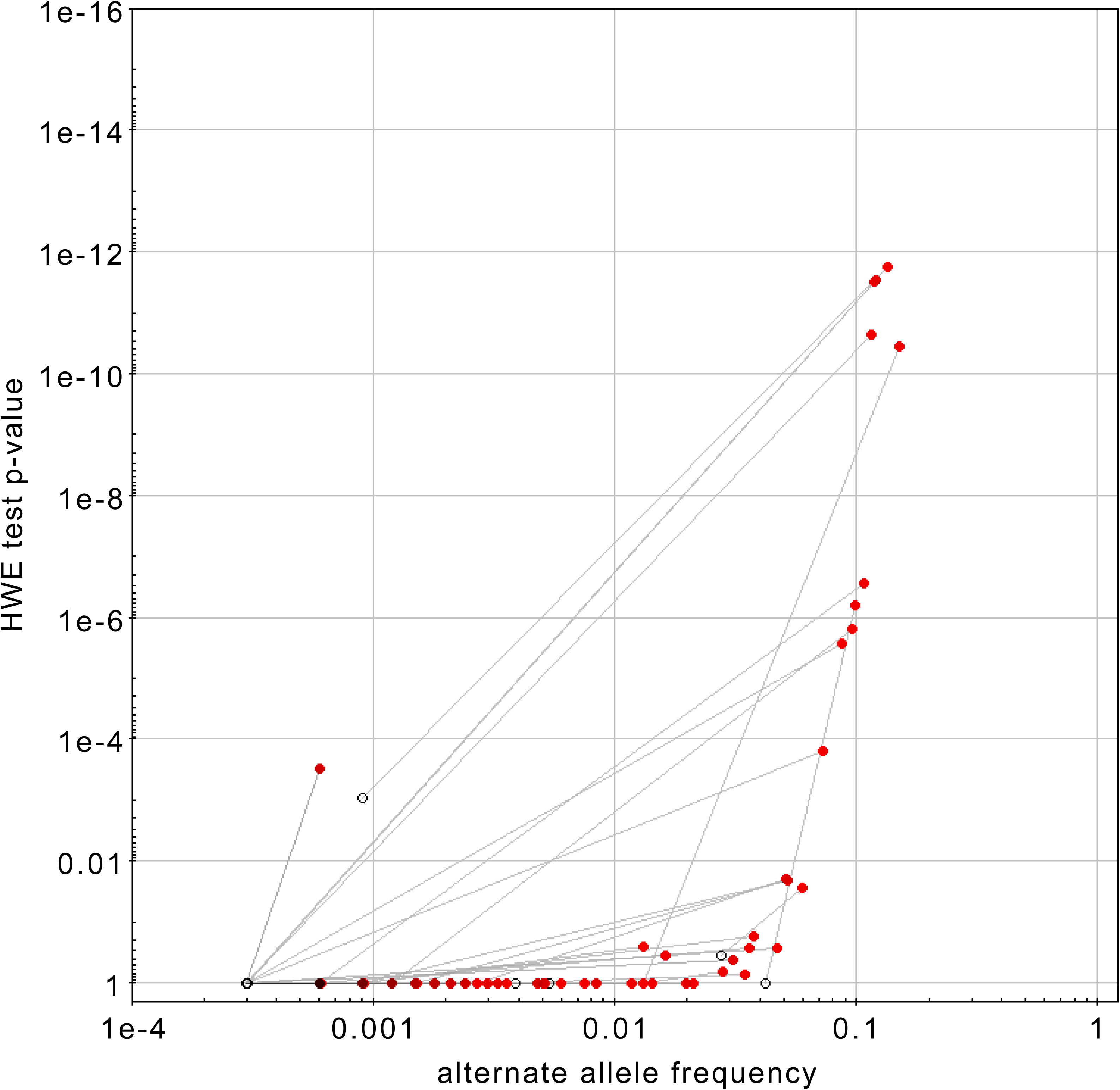
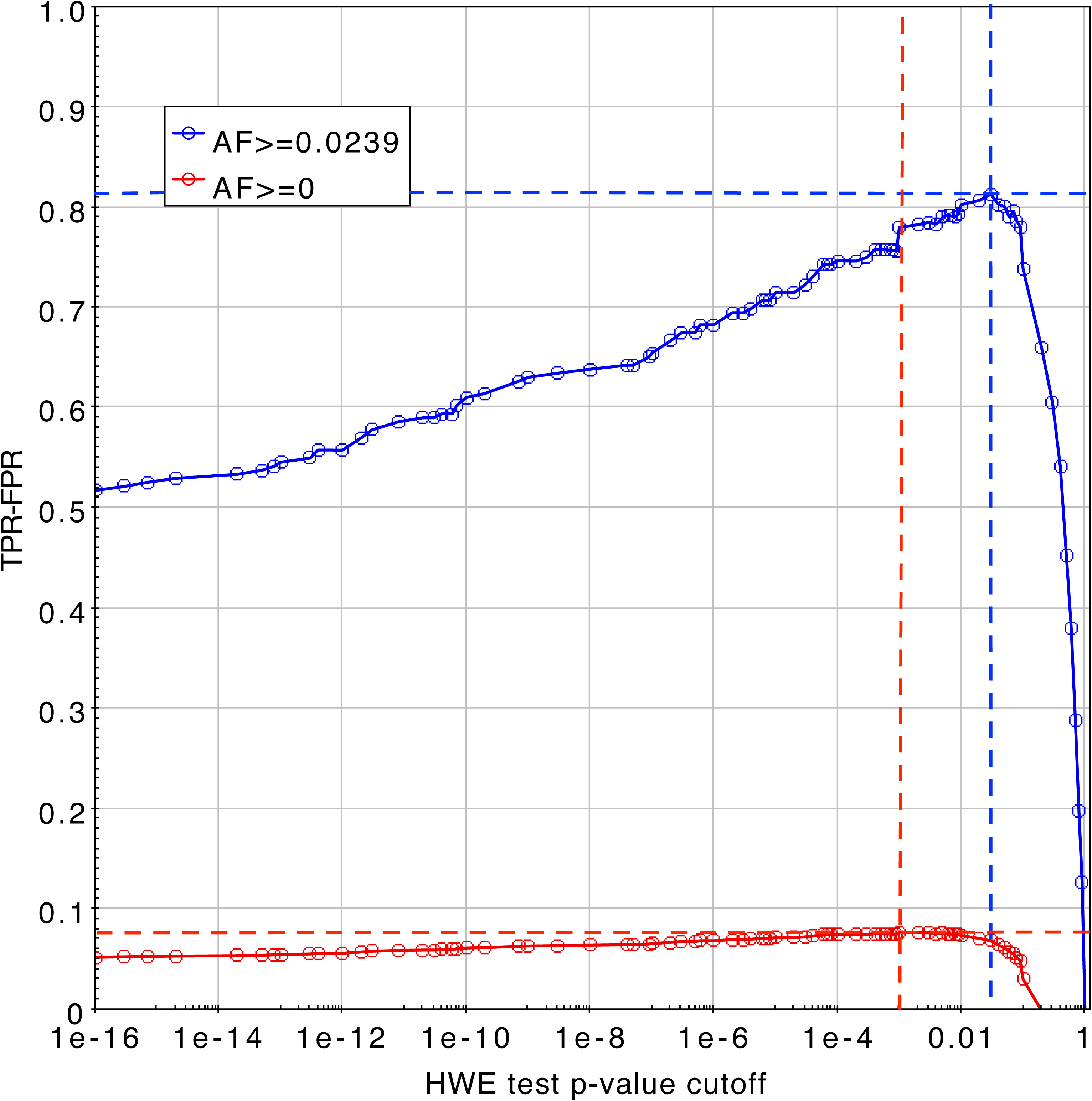
The difference between true positive rate (TPR) and false positive rate (FPR) as a function of HWE p-value cutoff. The cutoff is applied in the allele frequency range where the HWE test is effective (blue curve) and to the full allele frequency spectrum (red curve).

### Best practices of applying Hardy Weinberg Equilibrium test in the large scale variant calling quality control

In Text Box 1, we summarize the recommended site level quality control (QC) procedures to apply the HWE test in the variant calling with extremely large sample size.

##### Text Box 1 Best practice recommendation for the QC procedure

- Apply joint calling and genotyping across all samples (see Methods). (For variant callers which already make use of HWE information, we recommend keeping all sites and HWE p-value in the annotation as input of QC);
- Subset samples by population and in each population, apply the exact test of Hardy Weinberg Equilibrium at each variant site;
- Take a subset of samples from each population with shared samples in gold standard dataset (e.g. SNP array), recalculate HWE p-value for the subset of samples in each population, and identify true positive and false positive sites (see Discussion for the choice of golden data sets, and for cases when golden data set is not available);
- Based on the sample size in population *i*, find the limit allele frequency *f^i^*_c_ where HWE is informative at *f*>*f^i^*_c_ (see Methods);
- Find the optimal p-value cutoff *p^i^*_c_ which maximizes TPR-FPR at *f^i^*>*f^i^*_c_;
- Apply HWE p-value cutoff and filter variant sites with p-value *p^i^*<*p^i^*_c_ at *f^i^*>*f^i^*_c_ in the full call set in population *i*.

## Discussion

The effectiveness of HWE test for detecting sequencing artifacts in large sequencing cohorts is undocumented in previous literature, because of limitations on samples sizes. In this paper, we took a large cohort of whole genome sequencing samples with 6x-8x low coverage and revealed the limited power of HWE test at the rare end of the site frequency spectrum. As we showed as part of our results, the HWE test is very sensitive to the excess or deficit of heterozygotes samples at common false positive sites, but at the rare end (*f*<%), false positive sites with significant excess of heterozygotes samples cannot be distinguished from the true positives using the test. As was discussed in Results section, misalignment in local repetitive regions, false imputation and sequencing error, all plays a role in genotyping errors at the lower end of the frequency spectrum. We present a novel strategy to determine the cutoff on HWE test p-value, which effectively filters the false positive sites in the allele frequency range where HWE test is informative and achieves a better tradeoff between the sensitivity and specificity, compared to the filtering result with a constant p-value cutoff for all allele frequencies.

We recommend using high quality genotype data set as the quality control gold standard, which includes large number of samples commonly shared between the WGS call set. When selecting the golden data sets, we recommended considering following three factors. Firstly, the number of common samples in the golden data set should be large enough to avoid under-sampling of the rare false positive sites, and the samples should be within the same population so that the HWE p-value is not biased by any population stratification or significant population substructures. For instance, when the variant calling sample size is 10,000 and the HWE test limiting allele frequency is about 1% (Supplemental Figure 4), at least 500 samples in the same population should be present in the golden data set in order to assess the site quality with allele frequency at least an order of magnitude lower than the limiting allele frequency. Secondly, to assess the calling quality across the whole genome in a uniform manner, the golden data sets should adequately cover both the coding and non-coding regions, both the highly polymorphic sites and the rare variant sites. Using either the SNP array data containing only the common variants or the high coverage target sequencing call set (e.g. whole Exome sequencing, or WES) may not be sufficient. When large number of samples are available in quality control, combining different control data sets may provide balanced genomic coverage. Thirdly, the golden data set should include only the sites with high quality genotypes. The allele frequency dependent genotyping error in the golden data set will introduce noise in the quality assessment, and consequently bias the determination of HWE p-value cutoff. In case of SNP array data, using a consensus of different arrays can effectively avoid the array specific genotyping bias. In particular, any site with missing genotype in golden data set should be excluded from the negative controls, while sites with large fraction (e.g. 5%) of samples missing genotype, should be excluded from the positive controls. If the high coverage WES variant calling result is chosen as the golden data set, stringent site level genotype quality filter must be first applied. This is because in WES variant calling, the allele frequency dependency of genotyping error is not negligible and rare false positives may still be present due to the misalignment (see Supplemental Figure 3,6-7).

If the golden data set with shared samples is not available, we still recommend estimating the limit allele frequency *f*_c_ according to the sample size, and applying the HWE test to the sites with allele frequency *f*>*f*_c_. In this case, an empirical p-value cutoff needs to be determined from the optimal p-value cutoff with similar sample size in other large scale projects, but the tradeoff between sensitivity and specificity may not be optimal.

Since the HWE test has limited power in filtering the rare false positive variant sites, additional orthogonal filters may be used to improve the quality control in the rare end of the site frequency spectrum. For example, false positives due to local repetitive sequences can be filtered using low complexity regions (LCR) filter [10], while the sites with higher switching errors in the imputation and phasing may cluster in recombination hot spots regions in the genome. Since these regional or loci-specific filters are generally independent of the allele frequency of variant sites, they provide an orthogonal diagnosis to identify part of the rare false positive sites. Furthermore, per sample coverage information, when available, may also provide independent information to filter rare false positives with insufficient read depth at sample level. Improving the genotyping accuracy in the low coverage variant calling by making use of position specific genotype likelihood prior in a known population also helps to reduce the false positive rate. Besides, sample level quality control can be useful to identify rare false positives contributed by some outlying samples with extremely low or high coverage, or sequencing mishaps. Ultimately, given the sample size, increasing the sequencing coverage is still the most effective way to reduce the false positive calls in the rare end of site frequency spectrum.

## Conclusions

As the sequencing sample size continues to grow in the high throughput sequencing era, more rare variants are being discovered and quality control procedure to effectively remove the rare false positives is becoming increasingly important. Using the whole genome sequencing variant call set with more than 5000 samples, we revealed the limited power of exact test of HWE in filtering the rare false positive variant sites with allele frequency less than 1%. By analyzing the genotype discordance, we discussed the different causes of false calling and genotyping error in the low coverage sequencing. Given the availability of gold standard genotype data, we propose a novel strategy to ascertain the allele frequency range where the HWE test is informative and to determine the p-value cutoff which optimizes the overall sensitivity and specificity in the entire allele frequency spectrum. This strategy outperforms the conventional HWE filtering approaches with only an empirical constant p-value cutoff regardless the sequencing sample size. This can serve as a practical consideration for the variant level quality controls in the upcoming sequencing projects with tens or hundreds of thousand of samples.

## Materials and Methods

### Sequencing samples and variant calling

We took 5297 whole genome sequenced samples from the CHARGE project (Cohort for Heart and Aging Research in Genomic Epidemiology [15]). 3396 samples are European Americans (EuAm) and 1901 African Americans (AfAm). The samples were sequenced using Illumina HiSeq 2000 with an average coverage 6x-8x. The alignment was done using BWA [16] integrated in the Mercury pipeline [17]. We called biallelic SNPs across all 5297 samples, taking an ensemble variant calling approach goSNAP [18], which employs four variant callers GATK-HaplotypeCaller [19, 20] with gVCF option, GATK-UnifiedGenotyper [19, 20], SNPTools [21] and GotCloud [22], each enforced in a joint calling mode. To ensure a high quality variant call set, we applied a consensus filtering and selected 72,945,834 variant sites which were called at least in 3 out of all 4 callers. The genotype likelihood of each sample at each variant site was calculated using BAM specific Binomial Mixture Model (BBMM) algorithm implemented in SNPTools [21]. Imputation and phasing was done using SNPTools reference panel independent imputation engine. After imputation, we obtain phased genotypes of 5297 samples at 72,762,406 biallelic variant sites (52,116,900 in AfAm samples and 46,201,314 in EuAm samples). The site frequency spectrum of both populations follows similar patterns as in 1000 Genomes Project [4] (Supplementary Figure 1). We use the call set of both populations to evaluate the performance of Hardy Weinberg Equilibrium test.

### Golden data set for quality assessment

We use CHARGE SNP array data as the golden data set to assess the low coverage whole genome variant calling quality. Among 5297 CHARGE WGS sequencing samples, there are 3533 samples with SNP array genotype, 1683 EuAm samples and 1850 AfAm samples. The total number of autosomal variant sites in SNP array data is 228,963 across all 3533 samples. To ensure the quality of control data set, we further remove the control sites according to following criteria. In each population, we remove the sites called in WGS samples but have more than 5% of samples missing genotype in the array. We also remove sites which are not called in WGS samples and have any sample with missing genotype in the array. After filtering. the number of positive and negative control sites is 107,339 and 99,034, respectively, in EuAm samples (132,344 and 75,163 in AfAm samples).

Alternatively, we compare the WGS variant calling quality to the WES variant call set, which includes 4612 samples in common with CHARGE WGS, 1782 AfAm samples and 2830 EuAm samples. The sequencing coverage of WES is 80-100x. The number of control sites in AfAm and EuAm samples is in Supplemental Table 1.

### The exact test of Hardy Weinberg Equilibrium

We apply the exact test of Hardy Weinberg Equilibrium [6] to 52,116,900 variant sites called in 1901 AfAm WGS samples, and 46,201,314 variant sites called in 3396 EuAm samples. The exact HWE test is also applied to the filtered control sites in a subset of 1850 AfAm samples and 1683 EuAm samples, which are shared between the WGS and SNP array. Besides the p-value of exact HWE test, in each subset of samples and at each biallelic variant site, we also extract the number of samples with three genotypes, homozygous Ref/Ref (*N*_rr_), heterozygous Ref/Alt (*N*_ra_) and homozygous Alt/Alt (*N*_aa_). The alternate allele frequency is derived as *f* = (*N*_ra_+2*N*_aa_)/((*N*_rr_+*N*_ra_+*N*_aa_)×2).

### Calculate the limiting allele frequency of HWE test

To determine the p-value cutoff in HWE test, we first calculate the limiting allele frequency *f*_c_, above which the HWE is effective, based on the number of samples *N* in the same population. We keep the number of Alt/Alt samples *N*_aa_=0, and increase the number of heterozygous samples *N*_ra_ from 0, and *N*_rr_ = *N* – *N*_ra_. For each configuration of (*N*_rr_,*N*_ra_,*N*_aa_), we apply the exact test of HWE. When *N*_ra_ is small, the HWE *p*-value is always *p*=1. The HWE test is informative when the p-value starts to be less than 1, where we take number of heterozygotes *N*’_ra_ of this configuration and calculate the limiting allele frequency as *f*_c_ =*N*’_ra_ / 2*N*. We apply the HWE test only to variant sites with alternate allele frequency *f*>*f*_c_.

### Determine optimal p-value cutoff in the HWE test

By comparing to the golden data set among a subset of shared samples, we classify variant sites in WGS call set as true positive (shared in golden data set) and false positive sites (unique in WGS call set). For *T* true positive sites and *F* false positive sites with alternate allele frequency (derived based on genotypes in the WGS call set) higher than the limit allele frequency (*f* > *f*_c_), we apply the exact test of HWE and calculate the p-value. By changing p-value cutoff *p*_c_, we obtain *T*’ (*T*’<*T*) true positive sites and *F*’ (*F*’<*F*) false positive sites with *p*>*p*_c_. We calculate the true positive rate TPR(*p*_c_)=*T*’/*T* and false positive rate FPR(*p*_c_)=*F*’/*F*. The optimal HWE p-value cutoff is determined when TPR-FPR is maximized (Figure 4).

## Data

CHARGE dataset is accessible on dbGap with following study accession numbers, FHS: phs000651, ARIC: phs000668, CHS: phs000667. The permission is required to access the data. Participant consent is not required to access the data.

## Additional Files

Additional hwe.submit.supp.docs

